# The Fraser complex interconnects tissue layers to support basal epidermis and osteoblast integrated morphogenesis underlying fin skeletal patterning

**DOI:** 10.1101/2023.07.08.548238

**Authors:** Amy E. Robbins, Samuel G. Horst, Victor M. Lewis, Scott Stewart, Kryn Stankunas

## Abstract

Fraser Syndrome is a rare, multisystemic autosomal recessive disorder characterized by disrupted epithelial-mesenchymal associations upon loss of Fraser Complex genes. Disease manifestation and affected organs are highly variable. Digit malformations such as syndactyly are common but of unclear developmental origins. We explored if zebrafish *fraser extracellular matrix complex subunit 1 (fras1)* mutants model Fraser Syndrome-associated appendicular skeleton patterning defects. Approximately 10% of *fras1* mutants survive to adulthood, displaying striking and varied fin abnormalities, including endochondral bone fusions, ectopic cartilage, and disrupted caudal fin symmetry. The fins of surviving *fras1* mutants frequently have fewer and unbranched bony rays. *fras1* mutant fins regenerate to their original size but with exacerbated ray branching and fin symmetry defects. Single cell RNA-Seq analysis, *in situ* hybridizations, and antibody staining show specific Fraser complex expression in the basal epidermis during regenerative outgrowth. Fras1 and Fraser Complex component Frem2 accumulate along the basal side of distal-most basal epidermal cells. Greatly reduced and mislocalized Frem2 accompanies loss of Fras1 in *fras1* mutants. The Sonic hedgehog signaling between distal basal epidermis and adjacent mesenchymal pre-osteoblasts that promotes ray branching persists upon Fraser Complex loss. However, *fras1* mutant regenerating fins exhibit extensive sub-epidermal blistering associated with a disorganized basal epidermis and adjacent pre-osteoblasts. We propose Fraser Complex-supported tissue layer adhesion enables robust integrated tissue morphogenesis involving the basal epidermis and osteoblasts. Further, we establish zebrafish fin development and regeneration as an accessible model to explore mechanisms of Fraser Syndrome-associated digit defects and Fraser Complex function at epithelial-mesenchymal interfaces.

## INTRODUCTION

Fraser Syndrome is a rare autosomal recessive disorder characterized by a wide and varied range abnormalities across organ systems within (e.g., fluctuating asymmetry between left and right limbs) and between affected individuals. Limb and digit abnormalities, including cutaneous syndactyly (webbed digits), clinodactyly/camptodactyly (curved or bent digits, respectively), brachydactyly (short digits), and polydactyly (extra digits) are common (Slavotinek & Tifft, 2002; van Haelst et al., 2008; Pasu et al., 2011; Butt et al., 2014; Mahadevan et al., 2002). The hands and feet of individuals with Fraser Syndrome are as unique as the individuals themselves, reflecting the exceptional variability in penetrance and expressivity of Fraser Syndrome alleles.

Teleost fins represent evolutionary precursors of mammalian limbs (Coates, 1994; Shubin et al., 2006). Even fin rays (lepidotrichia), while dermal bone, share a deeply conserved gene regulatory network with tetrapod digit endochondral elements (Nakamura et al., 2016; Woltering et al., 2014). Therefore, zebrafish fins provide a particularly tractable model to study core mechanisms of digit skeletal patterning relevant to human disease. The zebrafish caudal fin and its rays is particularly well-studied, including as a leading model of robust, epimorphic regeneration (reviewed in Sehring and Weidinger, 2020). The caudal fin typically has eighteen principal rays, of which the central sixteen branch during development and regeneration (Arratia et al., 2008; Desvignes et al., 2022). Individual rays comprise two calcified and osteoblast-lined hemirays encasing intra-ray fibroblasts, nerves, vasculature and other cell types. Mesenchymal state pre-osteoblasts (pObs) proliferate within an outgrowth zone distal to each ray, then epithelialize to a bone-depositing differentiated state to extend forming rays. A laminin-containing basement membrane separates mature osteoblasts from a multi-layered epidermis that surrounds and interconnects the rays (Armstrong et al., 2017; Chen et al., 2015).

Sonic hedgehog (Shh) signaling from basal epidermal cells to adjacent pre-osteoblasts (pObs) promotes ray branching during development and regeneration (Zhang et al., 2012; Armstrong et al., 2017; Braunstein et al., 2021). Distally moving basal epidermal cells activate *sonic hedgehog a* (*shha)* expression while passing over pObs in the distal growth zone. The *shha*-expressing basal epidermal domain gradually splits during outgrowth, with *shha*-positive basal epidermal cells seemingly “pulling” pObs into divided pools to form a ray branch point. These signaling and potentially physical cell-cell interactions between epithelial (basal epidermal) and mesenchymal (pOb) state cells occur where the basement membrane remains immature. The contributions of this distinct extracellular environment to enable and/or modulate such cell interactions underlying ray branching morphogenesis are unresolved.

In Fraser Syndrome, disruptions to the basement membrane-localized Fraser Complex compromise epithelial-mesenchymal signaling, adhesion, and basement membrane rigidity (Smyth and Scambler, 2005). The Fraser Complex accumulates in the sublamina densa layer of the basement membrane separating epithelial from mesenchymal tissue layers. Mouse forward genetic screens identified pathogenic variants of Fraser Complex genes *Fraser extracellular matrix complex subunit* (*Fras1*) and *Fras1-related extracellular matrix protein 2* (*Frem2*), along with *Glutamate receptor interacting protein 1* (*Grip1*), a scaffold protein required for Fras1/Frem trafficking (Jadeja et al., 2005; McGregor et al., 2003; Takamiya et al., 2004; Vrontou et al., 2003). Loss of *Fras1* in mice results in subepidermal blistering of embryonic limb buds and digit syndactyly (Hines et al., 2016; Miller et al., 2013; Vrontou et al., 2003). Fraser Complex proteins are highly expressed in the embryonic zebrafish caudal fin (Dalezios et al., 2007; Gautier et al., 2008; Kiyozumi et al., 2006). Blistering is observed in the zebrafish caudal fin fold of *fras1* and other Fraser Syndrome-related mutant larvae, which also recapitulate characteristic Fraser Syndrome craniofacial abnormalities (Carney et al., 2010; Talbot et al., 2012; Talbot et al., 2016). However, adult phenotypes including any affecting fins have not been examined due to frequent larval lethality.

We explored Fraser Complex roles during fin formation using zebrafish *fras1^te262^* loss-of-function mutants (Carney et al., 2010; Talbot et al., 2016, 2012). We found ∼10% of *fras1* homozygous mutants survived to adulthood. All fins of surviving *fras1* mutant adults had varied skeletal defects reminiscent of tetrapod limb syndactyly, including intermittent ray branching, ray fusions, and overall fewer rays. Focusing on the caudal fin, we found abnormalities extended to the endochondral skeleton, including poor articulation of hypurals and radials and ectopic cartilaginous structures. *fras1* mutants regenerated amputated caudal fins to their original length. However, the regenerated fins acquired additional skeletal defects including further disrupted ray branching. Single-cell RNA-seq and *in situ* expression studies localized Fraser Complex transcripts and proteins to the distal-most basal epidermis and its mesenchyme-facing, basal side. Shh signaling between basal epidermis and pre-osteoblasts that promotes ray branching was retained in *fras1* mutant regenerating fins. However, the Fraser Complex did not assemble and extensive sub-epidermal blistering disrupted the normally tight association between the basal epidermis epithelium and underlying blastema mesenchyme, including pre-osteoblasts. We propose Fraser Complex is upregulated in distal outgrowing fins to establish a robust environment supporting interactions between basal epidermis and pre-osteoblasts that underlie skeletal patterning. Zebrafish *fras1* mutant fins and their regeneration provide a compelling context to explore Fraser complex functions at epithelial-mesenchymal interfaces during tissue morphogenesis and uncover deeply conserved mechanisms that may underlie Fraser Syndrome-associated syndactyly.

## RESULTS

### Fras1 promotes robust fin skeletal patterning including ray branching

The *fras1^te262^* mutant allele is reported as lethal around 10-12 days-post-fertilization (dpf) (Carney et al., 2010). However, we found 10.4% (*n* = 110/1057) of homozygous *fras1^te262^*mutants (hereafter referred to as *fras1^−/−^)* survived to adulthood under careful rearing conditions. Surviving adults required no special care and lived well over a year in standard housing conditions, albeit with reduced breeding efficiency. *fras1^−/−^* adults exhibited varied phenotypes, such as a protruding lower jaw and fin abnormalities including uneven pigment striping particularly evident in the caudal fin (Fig. 1A). Both paired (pectoral and pelvic) and median (dorsal, anal, caudal) fins showed extensive but highly varied fin and ray skeletal patterning defects, including asymmetrical fin shapes, reduced ray number associated with smaller fins, curved or warped rays, and decreased ray branching (Fig. 1A, Fig. 1 Supp. 1). Ray number differed between matched paired fins (i.e, pectoral fins) of the same fish (Fig. 1 Supp. 1C, D), consistent with fluctuating left-right asymmetry associated with human Fraser Syndrome.

**Figure 1.**
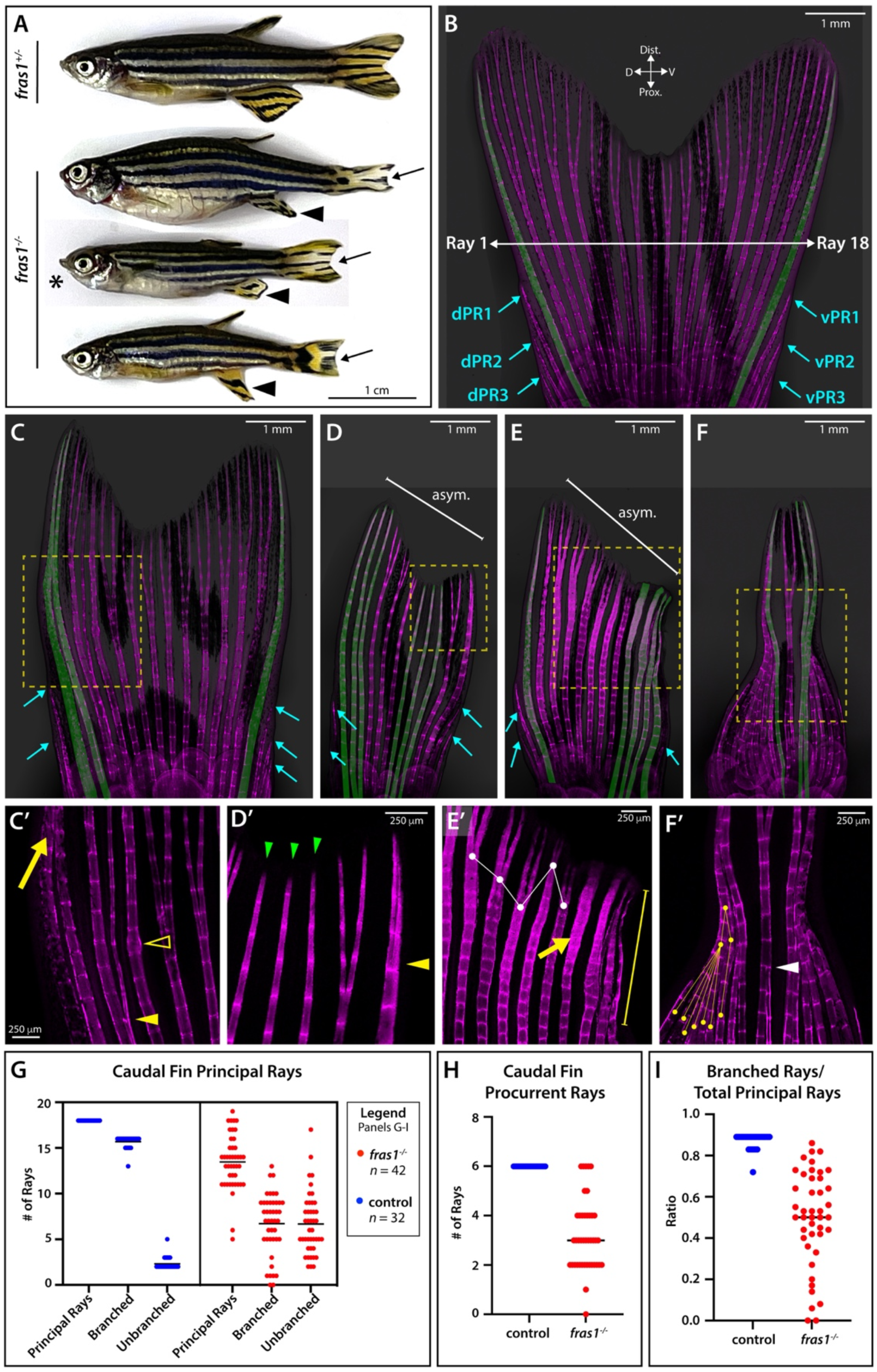
Fras1 promotes robust patterning of the caudal fin skeleton including the number of bony rays and ray branching. **(A)** Whole-mount imaged adult zebrafish representing three homozygous *fras1^−/−^*mutants and a heterozygous sibling. *fras1^−/−^* fish are variably runted with gross abnormalities including caudal fin pigment pattern defects (black arrows), misshapen anal fins (black arrowheads), and lower jaw protrusions (black asterisk). **(B-F)** Representative Alizarin Red-stained caudal fins of **(B)** control and **(C-F)** *fras1^−/−^* adult siblings. Unbranched rays are shaded green as a visual aid. Dorsal and ventral procurrent rays (dPR and vPR) are labeled with cyan arrows. Asymmetric fins **(D, E)** are indicated by white “asym.” brackets. **(C’-F’)** Various skeletal patterning abnormalities highlighted by zoomed in regions shown by yellow dashed boxes in **(C-F)** and noted below. **(C’)** Fused peripheral rays (yellow arrow), ectopic branching (yellow arrowhead), and fracture scar (yellow open arrowhead). **(D’)** Three unbranched rays (green arrowheads) followed by a branched ray followed by an ectopic, partially branched peripheral ray (yellow arrowhead). **(E’)** Uneven branch points (white connected dots), poorly articulated joints (yellow arrow), wavy misaligned rays (yellow bracket). **(F’)** Numerous fused rays (yellow connected dots) and isolated ray fusion (white arrowhead). **(G-I)** Individual data point plots showing control and *fras1^−/−^* adult caudal fin **(G)** principal ray number, **(H)** procurrent ray number, and **(I)** fin ray branching expressed as a fraction of total principal rays. Means are shown.

We focused on the caudal fin as an accessible model to further study Fraser complex roles in fin and fin ray development. Wildtype caudal fins typically have 18 principal, full length rays with 9 rays each symmetrically arranged on dorsal and ventral fin lobes. At least three shorter procurrent rays, sometimes called rudimentary rays, lie peripherally on each side of the peripheral rays. The inner 16 principal rays branch at least once, while the peripheral principal rays 1 and 18 remain unbranched. (Fig. 1B; Arratia et al., 2008; Coates, 1994). The caudal fin skeleton of *fras1^−/−^* fish (Fig. 1C-F) showed a range of abnormalities (Fig. 1B). *fras1^−/−^* caudal fins developed with fewer principal (count 5-18, *n =* 38) and procurrent rays (count 2-6, *n=* 37). Ray branching varied from normal (all central 16 of 18 rays) to completely absent (Fig. 1G-I). Additional abnormalities included overall fin asymmetry (Fig. 1D, E), ray fusions (Fig. 1C’, F’), precocious branching (Fig. 1C’), ectopic branching of peripheral principal rays (Fig. 1D’), unevenly positioned branch points, misaligned and wavy hemi-rays, and poorly articulated joints (Fig. 1E’). Therefore, Fras1 is required for fin skeletal patterning with variable expressivity and incomplete penetrance reminiscent of distal limb defects in humans with Fraser Syndrome.

### Fras1 contributes to endochondral and dermal bone patterning during fin development

Fin rays are dermal bone, unlike the fully endochondral skeleton of extant tetrapod limbs (Arratia et al., 2008; Coates, 1994). Therefore, we examined if zebrafish *fras1* also was required for endochondral fin skeletal pattering. We visualized the developing caudal skeleton of *fras1^−/−^* mutants by Alcian Blue and Alizarin Red staining of cartilage and calcified bone, respectively, at late larval stages. A representative heterozygote sibling control at 22 dpf (out of *n =* 7) (Fig. 2A) had five clearly defined hypurals centered around a hypural diastema (the gap between hypurals 2 and 3 around which principal rays symmetrically emerge; Desvignes et al., 2018; Desvignes et al, 2022). Age-(22 dpf, Fig. 2C) and size-matched (28 dpf, Fig. 2E, G) *fras1^−/−^* mutants showed varied caudal skeletal abnormalities including fusions between endochondral elements (Fig. 2C, E) and ectopic cartilaginous structures (Fig. 2C, E, G). Further, *fras1^−/−^* fish often displayed an extra dorsal hypural despite having fewer fin rays (Fig. 2C, E). As forementioned, while caudal fin rays normally are evenly spaced and segmented by joints (Fig. 1A), *fras1^−/−^* rays frequently were unaligned (Fig. 2C, G) and had poorly articulated joints (Fig. 2E, G). The hypural diastema was missing. The anal and dorsal median fins also showed endochondral skeletal defects. Control siblings had evenly spaced radials (Fig. 2B) whereas *fras1^−/−^* radials were variably truncated and/or contorted (Fig. 2D, F, H). The dorsal fin frequently had supernumerary posterior radials. We conclude Fras1 supports patterning of both endochondral and dermal bone fin skeletal elements.

**Figure 2.**
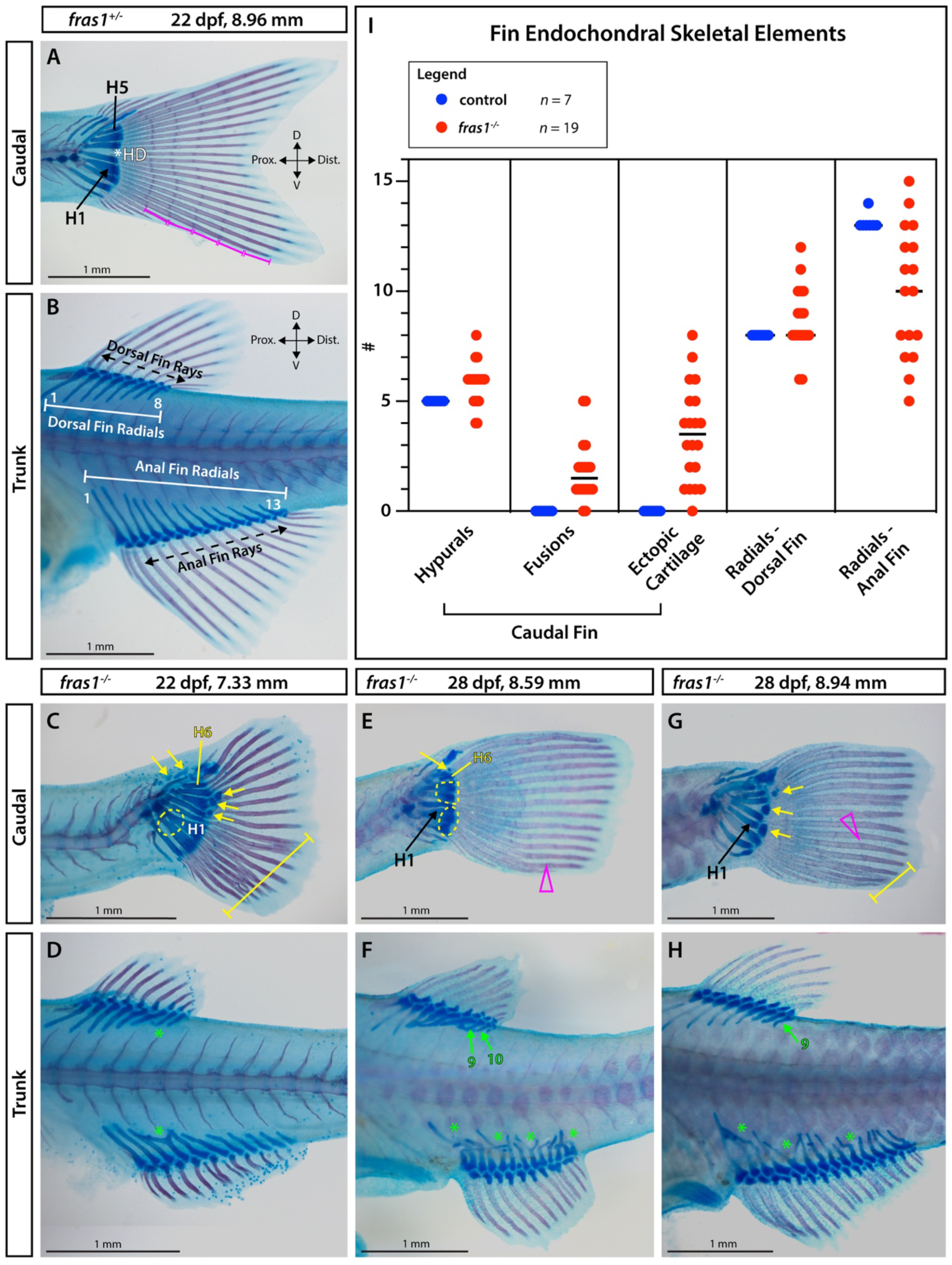
Fras1 contributes to both dermal and endochondral fin skeletal patterning. **(A-H)** Representative whole mount imaging of Alcian Blue (cartilage) and Alizarin Red (calcified bone) stained zebrafish of indicated *fras1* genotypes and ages. **(A)** Late larval *fras1^+/−^* control caudal fin. H1, hypural 1 and H5, hypural 5. Magenta brackets denote joint segmentation. **(B)** Trunk region of the same control larvae. Dorsal and anal fin radials are marked by white brackets and rays are marked by black dashed arrows. **(C-H)** Images of sibling *fras1^−/−^* late larvae age-matched **(C, D)** or size-matched **(E-H)** to control for frequent developmental delays, with highlighted defects described below. **(C, E, G)** Caudal fin endoskeletons (blue) are disorganized, often with a sixth hypural (H6, yellow), fused elements (yellow dashed lines), and ectopic cartilaginous structures (yellow arrows). **(E)** Caudal fin dermal rays (magenta) are variably misaligned (**C, G**; yellow brackets) and **(E, G)** have delayed joint development (magenta open arrowheads) even when size-matched to **(A)** younger controls. **(D, F, H)** Dorsal and anal fin endoskeletons (blue) have variably mispatterned radials (green asterisks), with dorsal fins often having extra radials (green arrows). Scale bars are 1 mm. **(I)** Data point plots scoring indicated fin skeletal elements or defects in *fras1^−/−^* mutants and control siblings. Means are shown.

### Fras1 stabilizes fin ray pattering without contributing to outgrowth during fin regeneration

Rapid and robust adult fin regeneration enables studying active roles of Fras1 during fin formation spanning wound healing through bone maturation. We tracked heterozygote control (*n* = 6) and *fras1^−/−^* (*n* = 9) siblings throughout caudal fin regeneration, imaging fins at uninjured, 3, 7, and 28 days-post-amputation (dpa) (Fig. 3). Control caudal fins progressively and robustly regenerated to their original size and bilobed shape with long peripheral rays, shorter central rays, and a branched ray field (Fig. 3B-D). *fras1^−/−^* fins at 3 dpa exhibited blistering between distal ray and epidermal boundaries (Fig. 3F, F’, N, N’) and excess distal epidermis (Fig 3F’, J’, N’). The outwardly apparent blisters seen at 3 dpa receded by 7 dpa (Fig. 3G, G’, K, K’, O, O’). By 28 dpa, *fras1^−/−^* caudal fins (*n* = 9/9) restored the same extent of fin tissue as controls (control ray length recovery from uninjured to 28 dpa for longest ray +/−5%; *fras1^−/−^*longest ray recovery +/−4%; shortest ray length recovery +/−8% for both groups) (Fig. 3S) but with exacerbated skeletal patterning defects including an altered number of principal fin rays (*n* = 8/9; Fig. 3I, L, M, P) and a further reduced number of branched rays (*n* = 9/9, paired t-test P=0.012; Fig. 3E, H, I, L, M, P, R).

**Figure 3.**
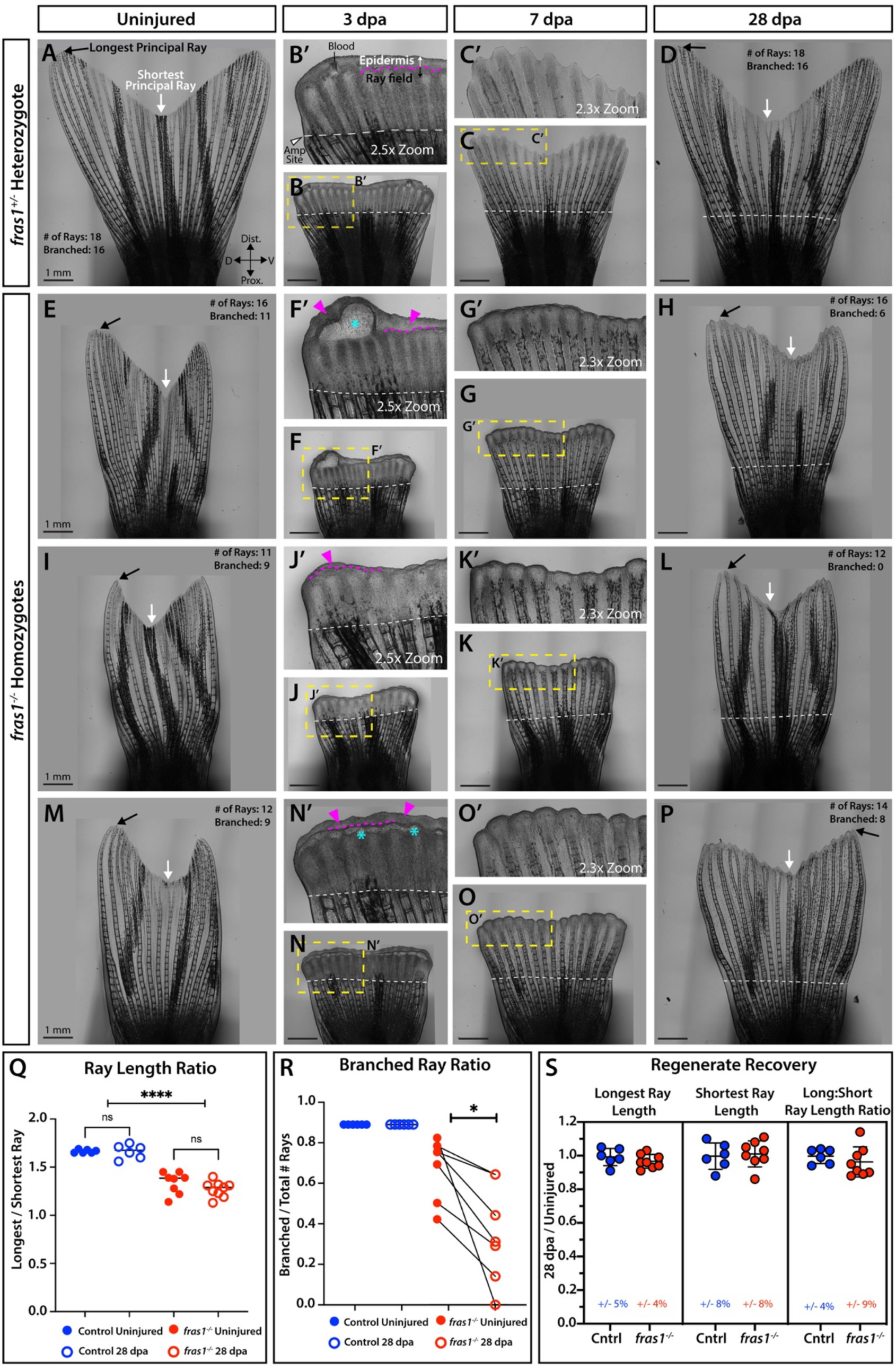
Fras1 robustly restores skeletal pattern including ray branching during adult caudal fin regeneration. **(A-D)** Images of representative *fras1^+/−^* heterozygote control adult caudal fins before injury (A, uninjured) and at 3 **(B)**, 7 **(C)**, and 28 **(D)** days post amputation (dpa). **(B’, C’)** Zoom insets of the yellow boxed regions in B and C. **(B’)** Black arrows **(A)** indicate the longest principal ray and white arrows indicate the shortest. Magenta dashed line denotes the approximate boundary between the outgrowing ray field and distal epidermis. White dashed lines indicate the plane of amputation. **(E-P)** Three representative *fras1^−/−^* homozygote siblings at the same regenerative time points as **(A-D)** highlighting defects noted below. At 3 dpa, mutants exhibit blistering (cyan asterisk; **F’, N’**) and extended epidermal tissue distal to the ray field (magenta arrowheads; **F’, J’, N’**). **(G, G’, K, K’, O, O’)** Blistering subsides by 7 dpa. **(H-P)** Fully regenerated mutant fins have **(H)** fewer branched rays, **(L)** no branched rays, or more equal ray length ratios (i.e., less pronounced V-shape) **(C)** compared to their uninjured states. Scale bar lengths are indicated. **(Q-S)** Individual data point plots comparing fin shape as well as ray length and branching restoration upon regeneration between control and *fras1^−/−^* animals. **(Q)** Ray length ratios between the longest and shortest principal rays before injury (solid circles) and at 28 dpa (unfilled circles). Uninjured and regenerated ray length ratios are unchanged within genotypes (control P = 0.92, *fras1*^−/−^ P=0.22); paired t-tests) but are significantly different between genotypes (P <0.0001 for uninjured and regenerated fins; unpaired t-tests). **(R)** The fraction of branched over total principal rays, with connecting lines indicating the same fish assessed before injury and at 28 dpa. Ray branching frequency decreased after regeneration in all *fras1^−/−^*animals (P = 0.012; paired t-test) but no control animals (P = 1). **(S)** The fraction of regenerated to original ray length for the longest and shortest principal rays and the ratio of the two values. Means and SD are shown.

We also noted *fras1^−/−^* caudal fins exhibited a shallower bi-lobed “V” shape compared to control samples in both uninjured and regenerated states. Concordantly, the ratio of the longest to shortest ray was significantly lower in *fras1^−/−^* vs. control caudal fins in both uninjured and 28 dpa regenerated states (unpaired t-test P <0.0001 for both uninjured and 28 dpa) (Fig. 3E, I, M, Q). In contrast, ray proportions were unchanged between uninjured and 28 dpa control fish (paired t-test P = 0.92, *n* = 6) or uninjured vs. 28 dpa *fras1^−/−^* fish (paired t-test P = 0.22, *n* = 8). The longest ray switched from the dorsal-most (uninjured, Fig. 3M) to ventral-most principal ray after regeneration in one *fras1^−/−^* individual (*n* = 1/9; Fig. 3P). Taken together, the exacerbated and varied phenotypes in *fras1^−/−^* regenerated fins indicates Fras1 stabilizes tissue patterning during both developmental and regenerative fin outgrowth.

### Fraser Complex transcripts and proteins are enriched in distal fin regenerates and support pre-osteoblasts alignment

The subepidermal blistering in early *fras1^−/−^* fin regenerates characterizes Fraser syndrome-associated defects in other contexts. Therefore, zebrafish fin regeneration provides a new and compelling model to explore Fraser Complex function in modulating interactions between adjacent epithelial and mesenchymal cells. We examined Fraser Complex transcript and protein localization during fin regeneration, anticipating expression in cells adjacent to the basement membrane separating basal epidermis from blastema (Fig. 4A, B). We first mined our existing single-cell RNA-Seq dataset of 3 and 7 dpa regenerating fin cells, focusing on the unique clusters for superficial epidermis, basal epidermis, and combined mesenchyme, fibroblast and osteoblast-lineage cells (Fig. 4C; Lewis et al., 2023). As reported by bulk RNA-Seq analyses (Nauroy et al., 2019), Fraser Complex transcripts *frem2a, frem2b*, and *frem3* were highly expressed in the basal epidermal cluster, with *fras1* being the single most enriched basal epidermal marker in our dataset (Fig. 4D, Fig. 4 Supp. 1). The Fraser complex transcript *frem1a* was expressed sparsely in basal epidermal and mesenchyme/fibroblast/osteoblast clusters, and *frem1b* showed lower levels in the fibroblast/osteoblast cluster and scarce expression in superficial epidermis (Fig. 4D, Fig. 4 Supp. 1). These data are consistent with mouse studies showing Fraser Complex transcript expression in associated epithelial and mesenchymal tissue layers with the secreted proteins complexing in the intervening basement membrane (Jadeja et al., 2005; McGregor et al., 2003; Takamiya et al., 2004; Vrontou et al., 2003).

**Figure 4.**
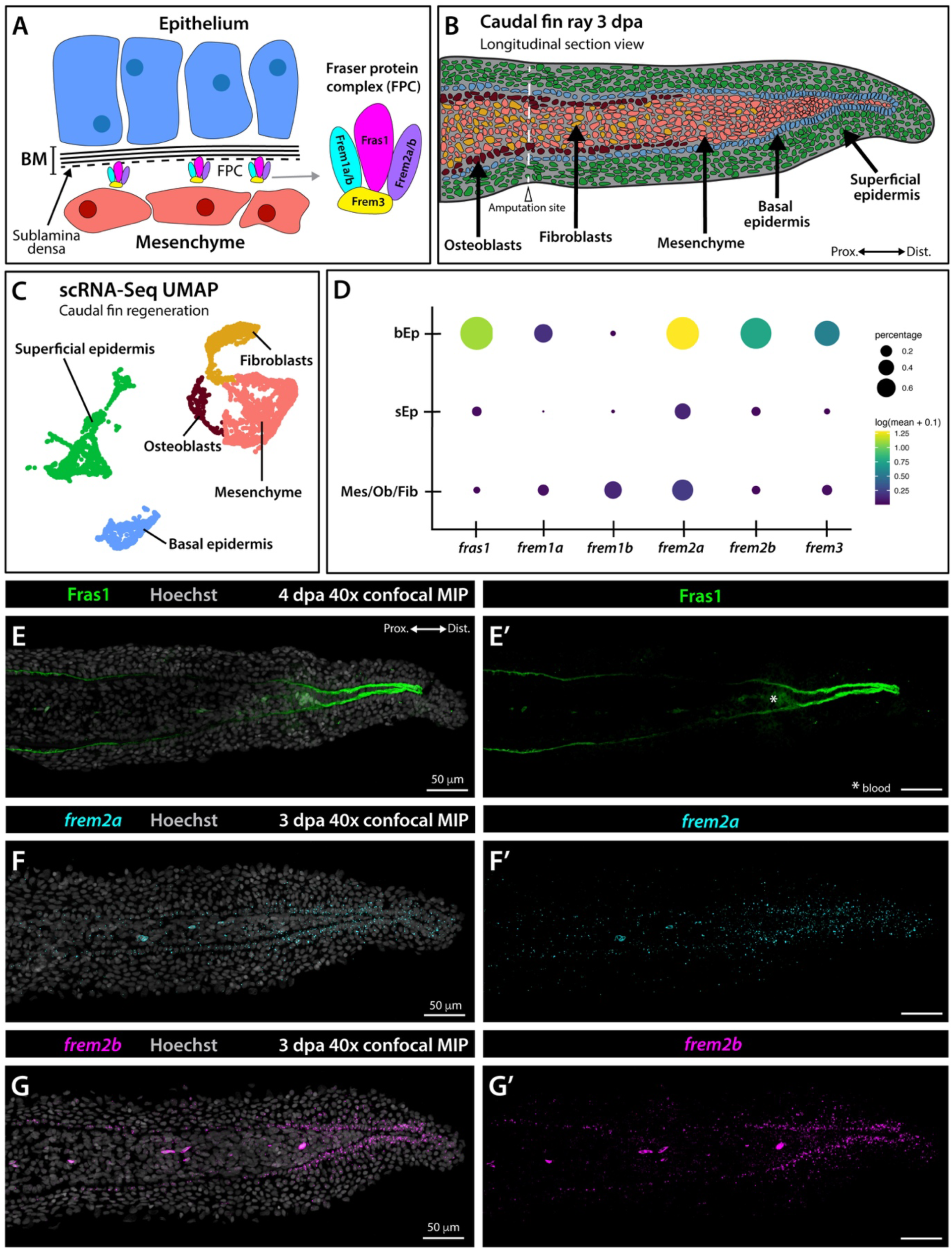
Fraser Complex transcripts and Fras1 protein are basal epidermal-expressed and distally-enriched in regenerating fins. **(A)** Schematic of epithelial (blue cells) and mesenchymal (pink cells) tissue layers separated by basement membrane (BM, black lines). The Fraser Complex (FPC, multicolored) localizes to the innermost BM layer (sublamina densa, dashed black line) and comprises Fras1 (magenta), Frem1a/b (cyan), Frem2a/b (purple), and Frem3 (yellow). **(B)** Longitudinal section schematic of tissue organization in a 3-day post amputation (dpa) regenerating caudal fin. **(C)** Single cell RNA-Seq clustering of regenerating 3 and 7 dpa zebrafish fin tissue shows unique clusters for basal epidermis (blue), superficial epidermis (green), and mixed mesenchyme/osteoblast/fibroblast lineages (dark red, pink, yellow) in UMAP space. **(D)** Bubble plot of FPC transcript expression in the basal epidermis, superficial epidermis (sEp), and mixed mesenchyme/osteoblast/fibroblast (Mes/Ob/Fib) clusters. **(E-G)** 40x confocal maximum intensity projections of regenerating fin longitudinal sections stained as follows: **(E)** Fras1 immunostaining (green) at 4 dpa, **(F)** ViewRNA fluorescent *in situ* hybridization for *frem2a* (cyan) at 3 dpa, and **(G)** ViewRNA fluorescent *in situ* hybridization for *frem2b* (magenta) at 3 dpa. Hoechst-stained nuclei are grey. **(E’-G’)** Single channel fluorescence images of **(E-G)**. Scale bars are 50 μm.

We antibody stained 4 dpa regenerating fin sections to define Fras1 protein spatial expression *in situ*. Fras1 was highly enriched in the distal-most tissue separating basal epidermis from blastemal mesenchyme. Fras1 was also present in proximal, re-formed tissue, in association with the mature basement membrane separating basal epidermis from bone-lining osteoblasts (Fig. 4E, E’; Armstrong et al., 2017; Chen et al., 2015). Fluorescent *in situ* hybridization demonstrated *frem2a* transcripts were highest in far distal basal epidermis with lower expression in blastemal mesenchyme and superficial epidermis. The *frem2b* pattern was similar but more specific to basal epidermal cells (Fig. 4F-G’). We conclude Fraser Complex transcripts are upregulated in distal basal epidermis of regenerating fins to produce locally accumulating Fraser Complex.

We examined expression of Fras1 relative to pre-osteoblasts in 4 dpa regenerating fin sections given skeletal defects in *fras1^−/−^* mutants. Fras1 protein mostly accumulated distal to blastema-lining Runx2-expressing pre-osteoblasts (Fig. 5A). Lower Fras1 was found lateral to the most distal pre-osteoblasts. Fras1 protein unsurprisingly was absent in *fras1^−/−^* mutants (*n* = 6) (Fig. 5B). Runx2+ pre-osteoblasts were clumped and multi-layered rather than evenly spaced along the basal epidermis (Fig. 5A, B). We also examined Frem2 protein localization relative to Runx2+ pre-osteoblasts. Like Fras1, Frem2 was highly enriched distally between the basal epidermis and blastema mesenchyme and largely distal to pre-osteoblasts (Fig. 5C, C’). Frem2 was also present extracellularly within the superficial epidermis layer enveloping distal basal epidermal cells (Fig. 5C”, *n* = 6). In *fras1^−/−^*mutants, Frem2 protein was almost completely absent (Fig. 5D, D’) and what protein remained appeared intracellularly retained within basal epidermal cells (Fig. 5D”, *n* = 6). We conclude Fras1 supports Frem2 secretion and assembly into Fraser Complex-containing basement membrane. Further, we find that Fras1 promotes pre-osteoblast alignment, albeit likely indirectly given the Fraser Complex minimally associated with pre-osteoblasts.

**Figure 5.**
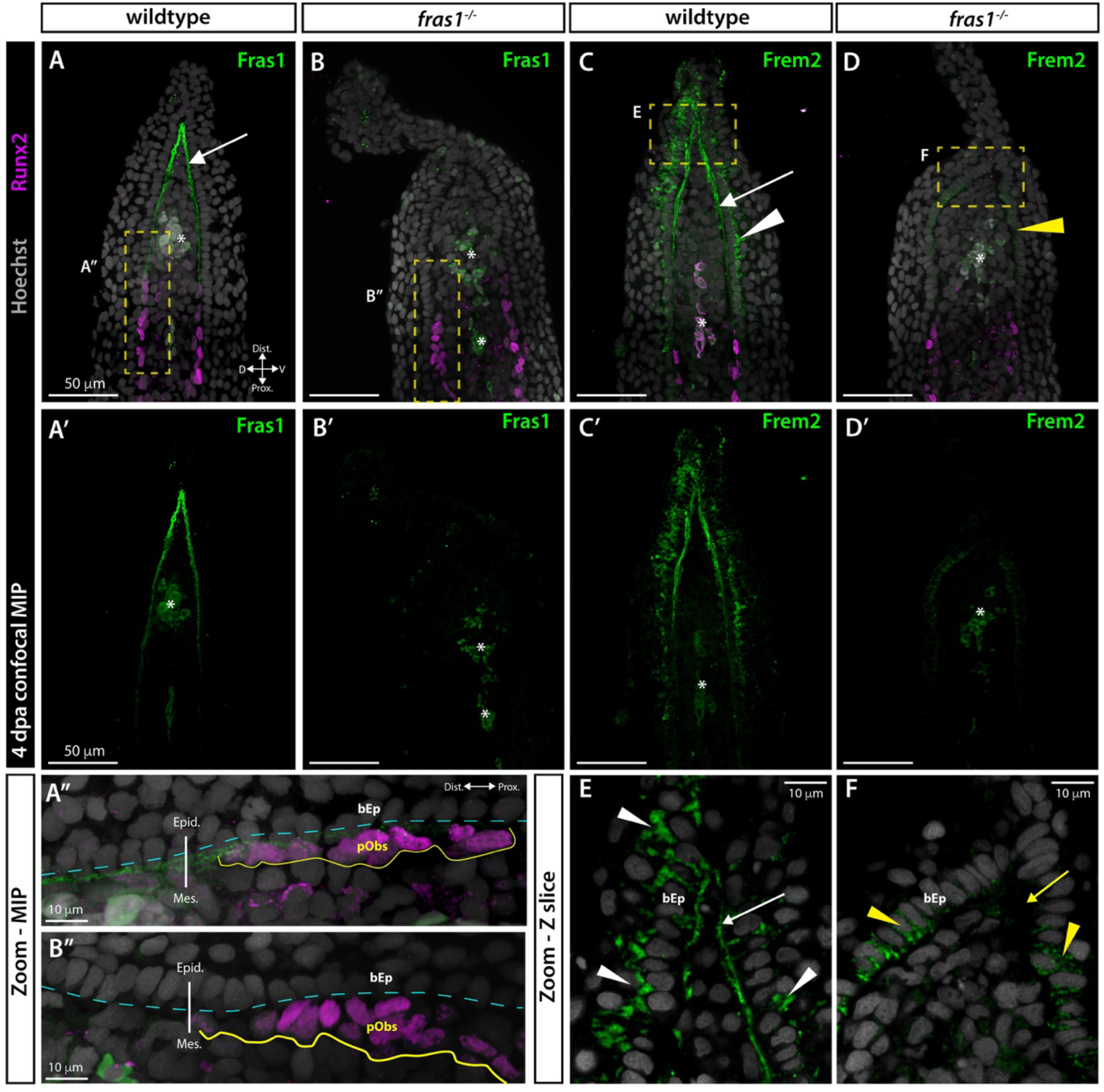
Fras1 promotes Fraser Complex assembly and pre-osteoblast organization during fin regeneration. **(A-D)** Confocal images of immunofluorescent stained paraffin sections of 4 day post amputation (4 dpa) regenerating caudal fins. Hoechst (gray) stains nuclei and anti-Runx2 antibody staining (magenta) marks pre-osteoblasts. Asterisks indicate blood autofluorescence. **(A-D)** Maximum intensity projection 3-channel overlays. **(A’-D’)** Single channel showing Fras1 or Frem2 immunostaining (green). **(A”, B”)** Maximum intensity projection zoomed views of the yellow boxed regions in **(A-B)**. **(E, F)** Single optical slice zooms of the yellow boxed regions in **(C-D)**. **(A, A’)** Fras1 (green) in wildtype fins is strongly expressed in the distal basement membrane with diminishing expression extending proximally. **(A”)** Fras1 is expressed at low levels between Runx2+ pre-osteoblasts (yellow outline) and epidermal cells. Epid, epidermis; Mes., mesenchyme. The cyan dashed line indicates the approximate basement membrane position. **(B-B”)** Fras1 (green) is lost and pre-osteoblasts are disorganized in *fras1^−/−^*mutants. **(C, C’, C”)** Frem2 (green) is expressed in the distal basement membrane (white arrow) and in the extracellular space around basal epidermal cells (bEp, white arrowheads, **C”**). **(D-D”)** Frem2 expression (green) in *fras1^−/−^* mutants is absent from distal basement membrane (yellow arrow) and the remaining protein appears intracellular within basal epidermal cells (yellow arrowheads). Scale bar lengths are indicated.

### Fras1-mediated skeletal patterning functions independently of Shh/Smo signaling

Fin ray branching requires basal epidermal Sonic hedgehog/Smoothened (Shh/Smo) signaling to divide underlying pre-osteoblast pools during development and regeneration (Armstrong et al., 2017; Braunstein et al., 2021; Cardeira-da-Silva et al., 2022; Quint et al., 2002). We considered if this epithelial-to-mesenchymal signaling was disrupted in *fras1^−/−^*mutants given frequent ray branching defects and disorganized Runx2+ pre-osteoblasts. *shha* is transcriptionally activated in distal basal epidermal cells as they collectively move distally and pass over pre-osteoblasts of the outgrowth zone (Zhang et al., 2012; Braunstein et al., 2021). *shha*-expressing basal epidermal domains progressively divide during fin outgrowth, likely driving the subsequent Shh/Smo-dependent splitting of pre-osteoblast pools. We assessed if *shha* expression during fin regeneration was Fras1-dependent using *Tg(−2.4shha:gfpABC)* (Ertzer et al., 2007; henceforth referred to as *shha:GFP*). *shha:GFP* was expressed in distal ray-associated domains in both heterozygote control and *fras1^−/−^* mutant 8 dpa regenerating caudal fins (*n* = 3 each). Likewise, each ray-associated *shha:GFP* domain of *fras1^−/−^*regenerating fins still split by 8 dpa, when ray branching is well underway, even though many *fras1^−/−^* fin rays did not branch (Fig. 6A, A’, C, C’). Therefore, Fraser complex is not required for *shha* induction nor the *shha*-expressing basal epidermal domain splitting underlying ray branching.

**Figure 6.**
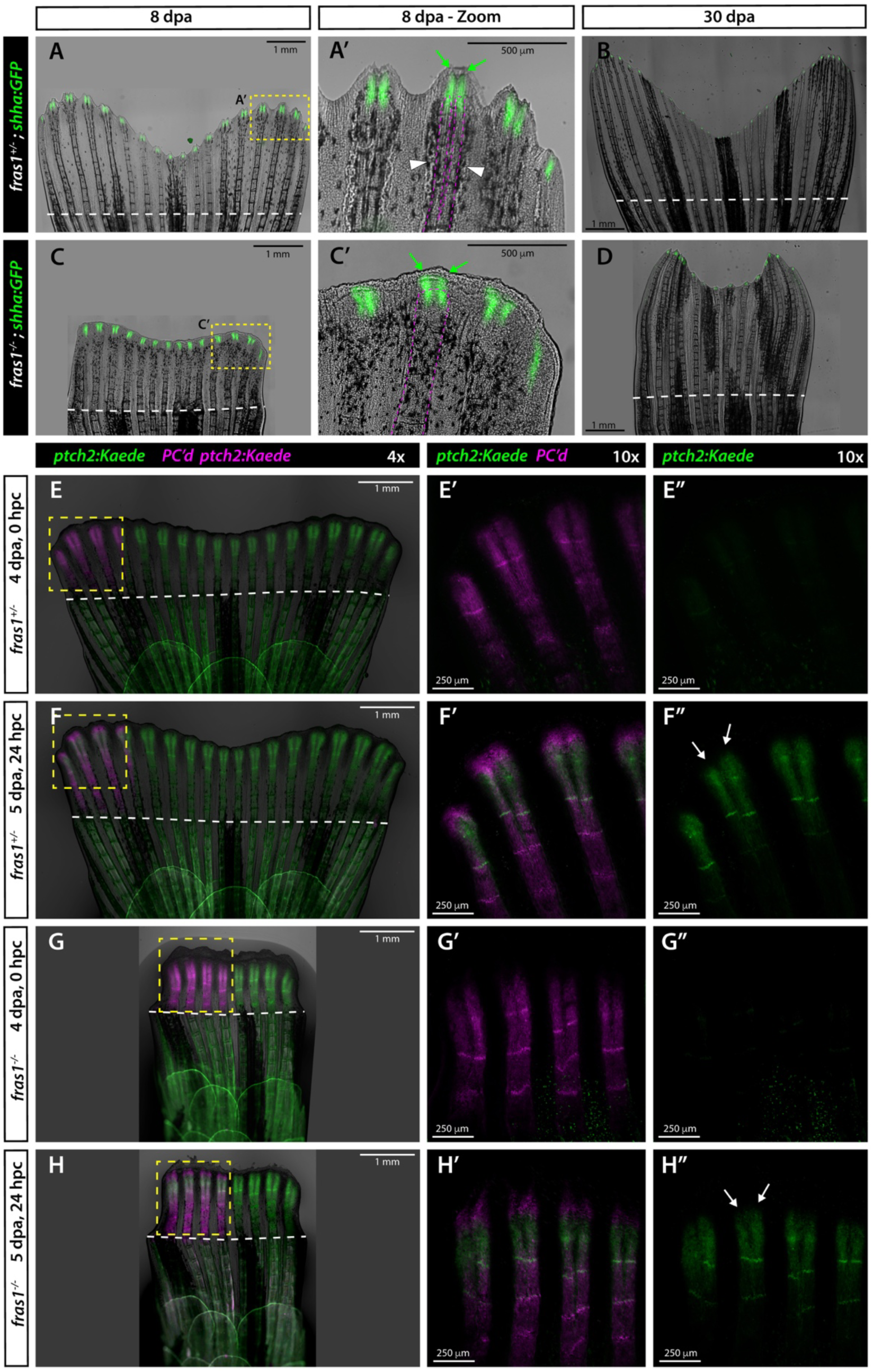
The Fraser Complex is not essential for Sonic hedgehog/Smoothened signal transduction during fin regeneration. **(A-D)** Whole mount images of regenerating caudal fins of **(A-B)** *fras1^+/−^* control sibling and **(C-D)** *fras1^−/−^* zebrafish carrying the *shha:GFP* reporter. **(A)** At 8 days post amputation (dpa), *shha:GFP* is expressed in basal epidermal cells overlying the distal outgrowth zone of each ray. **(A’)** Zoom of ventral ray regions showing GFP domain splitting (green arrows) associated with a branched ray (magenta dashed outline). **(C, C’)** A *fras1^−/−^* fish at 8 dpa with split distal GFP domains (**C’**; green arrows) even while the underlying rays fail to branch (magenta dashed outline). **(B, D)** Regeneration is complete at 30 dpa, with only residual distal GFP. The *fras1^−/−^*animal **(D)** fails to regenerate branched rays. White dashed lines indicate the amputation site. **(E-H)** Whole mount images of regenerating caudal fins from *ptch2:Kaede* sibling zebrafish of indicated *fras1* genotype. **(E, G)** 4 dpa fins imaged immediately after *ptch2:Kaede* is photoconverted from green to red (shown in magenta) fluorescence in the marked field (yellow dashed boxes, 0 hours-post-conversion hpc). **(F, H)** The same fins imaged 24 hours later (at 5 dpa, 24 hpc) showing newly produced green Kaede domains in both genotypes (**F”, H”**; white arrows). **(E’-H’)** Double fluorescent and **(E”-H”)** single fluorescence channels. Scale bar lengths are indicated.

We next considered if Fras1 facilitates Shh/Smo signal transduction to pre-osteoblasts. We photoconverted the well-established Shh/Smo activity reporter *Tg(ptch2:Kaede)* (Huang et al., 2012; henceforth *ptch2:Kaede*) in distal ray regions at 4 dpa from green to red (false colored magenta) (Fig. 6E, G). Both heterozygote controls and *fras1^−/−^* mutants (*n* = 6 each) produced new domains of green Kaede protein at 24 hours post-conversion, even in clearly abnormal *fras1^−/−^* fins (Fig. 6F, H). *ptch2:Kaede* photoconversion assays during fin development yielded similar results (Fig. 6 Supp. 1). We conclude Shh/Smo signal transduction in zebrafish fins does not require the Fraser Complex. However, the Fraser Complex at least supports the Shh/Smo downstream cell behaviors that split pre-osteoblasts given frequent ray branching defects in *fras1^−/−^*mutants. The varied phenotypes in *fras1^−/−^* mutants and predominant expression of the Fraser complex distal to the Shh/Smo signaling zone, including pre-osteoblasts, suggest an indirect and/or supporting role.

### Fras1 maintains epidermal-mesenchymal continuity in regenerating caudal fin rays

The morphological changes to *fras1^−/−^* regenerating rays (e.g., blistering, blunted ends, disorganized bone) led us to investigate how Fras1 contributes to tissue organization during caudal fin regeneration. We hematoxylin and eosin (H&E)-stained 4 dpa regenerating caudal fin sections of heterozygote control (*n* = 7) and *fras1^−/−^* animals (*n* = 8). The blastema mesenchyme of control fins was continuously associated with a surrounding single-cell layer of basal epidermis (bEps) further encased by superficial epidermis (sEp) (Fig. 7A). *fras1^−/−^* regenerates variably exhibited an uneven shape with acellular gaps and pockets (or, “blisters”) concentrated distally (Fig. 7B-D). Single confocal optical slices of eosin-stained fins (Fig. 7E-H) revealed fully penetrant but varied blistering with intra-epidermal acellular pockets and tissue separations between the basal epidermis and underlying blastema mesenchyme. We frequently also observed excess “flaps” of epidermal tissue distal to the blastema (Fig. 7F-H). We conclude Fras1 maintains tissue continuity in the distal ray outgrowth region of regenerating fins to enable robust restoration of skeletal pattern.

**Figure 7.**
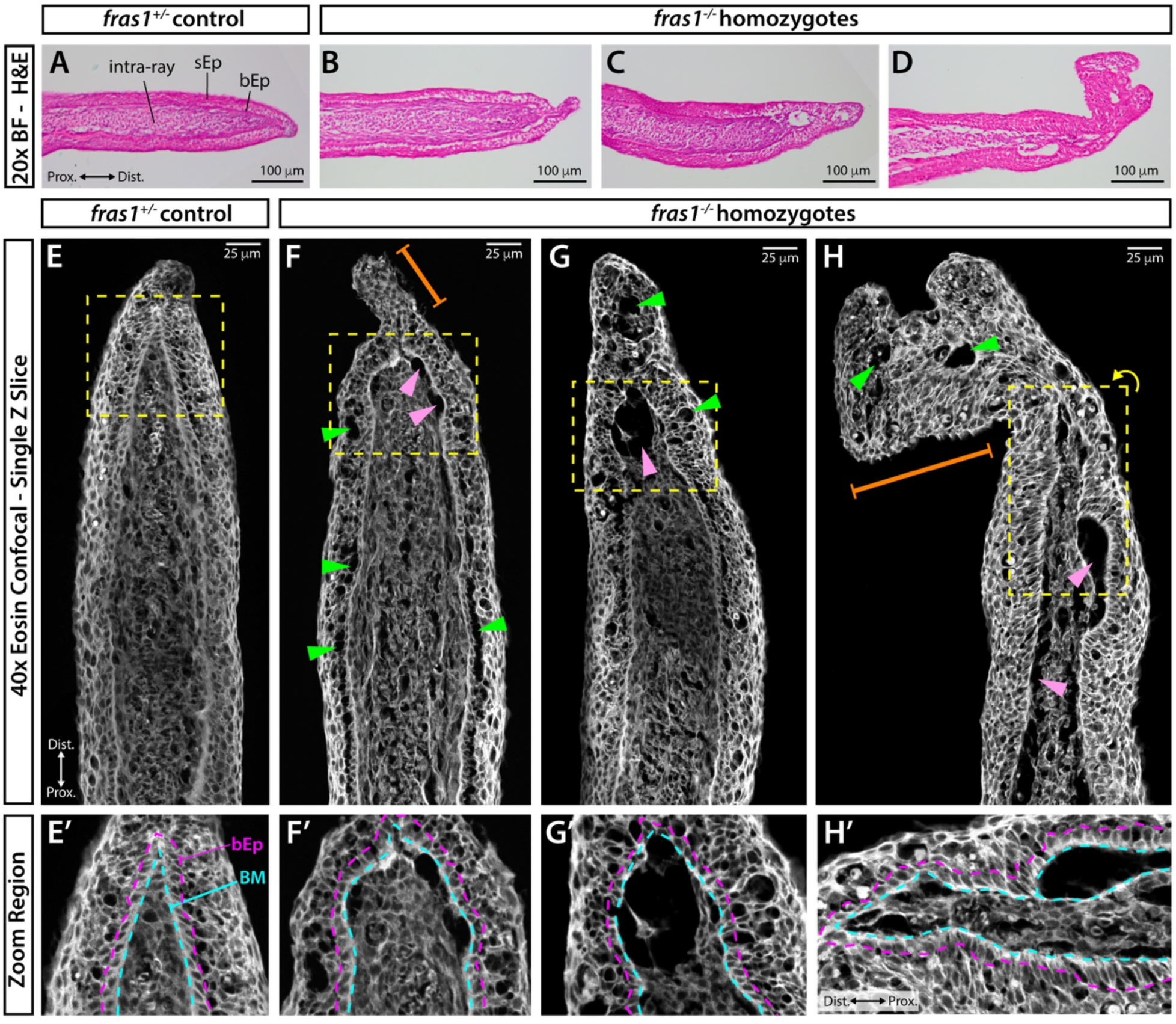
The Fraser complex maintains distal epithelial-mesenchymal tissue layering during fin regeneration. **(A-H’)** Hematoxylin and eosin-stained longitudinal sections of 4 day post amputation (4 dpa) regenerating caudal fins from *fras1^+/−^*control and *fras1^−/−^* adult zebrafish. **(A-D)** Brightfield images with hematoxylin-marked nuclei (purple) and eosin-marked cytoplasm and ECM (pink). Representative *fras^+/−^* control **(A)** and *fras1^−/−^* fish (**B-D**) are shown. sEp, superficial epidermis; bEp, basal epidermis; Mes., mesenchyme. **(E-H)** Confocal optical slices showing fluorescent eosin signal from the samples in **(A-D)**. Green arrowheads indicate epidermal blistering. Pink arrowheads indicate mesenchymal blistering. Orange brackets indicate excess epidermal tissue. **(E’-H’)** Zoom regions of yellow dashed boxes in **(E-H)** with approximate boundaries of the distal-most forming basement membrane (BM; cyan dashed line) and basal epidermis (bEp; magenta dashed line). Scale bars are indicated.

## DISCUSSION

We show that *fras1* loss-of-function zebrafish develop variably abnormal fins affecting both endochondral and dermal skeletons. These skeletal patterning defects, including supernumerary cartilages, fin asymmetry with aberrant ray numbers, and deficient ray branching resemble varied limb/digit abnormalities seen in Fraser Syndrome. The most common limb defect in Fraser Syndrome is cutaneous syndactyly (webbed digits, frequency 54%-79%; (Slavotinek and Tifft, 2002; van Haelst et al., 2008), a phenotype also observed in mouse models (Jadeja et al., 2005; McGregor et al., 2003; Miller et al., 2013; Smyth et al., 2004; Takamiya et al., 2004; Vrontou et al., 2003). Zebrafish Fraser complex mutants cannot model cutaneous syndactyly as fins and their component rays and endochondral bones are epidermally webbed. However, Fraser syndrome syndactyly also includes digit skeletal fusions and ectopic skeletal elements (Butt et al., 2014; Mahadevan et al., 2002; Pasu et al., 2011). Further, fin dermal bone rays and tetrapod limb digits may share deep evolutionary homology despite their substantial anatomical divergence (Boisvert et al., 2008; Freitas et al., 2012; Letelier et al., 2018; Nakamura et al., 2016; Sordlno et al., 1995). Therefore, fin endochondral and ray patterning defects in *fras1^te262^* homozygous zebrafish provide new models to understand origins of Fraser syndrome limb abnormalities affecting skeletal pattern (i.e., complex syndactyly, loss of digits). Centrally, our results indicate the Fraser complex establishes an environment supporting cooperative epidermal and adjacent mesenchymal, including pre-osteoblast, interactions for robust fin skeletal patterning (Fig. 8).

**Figure 8.**
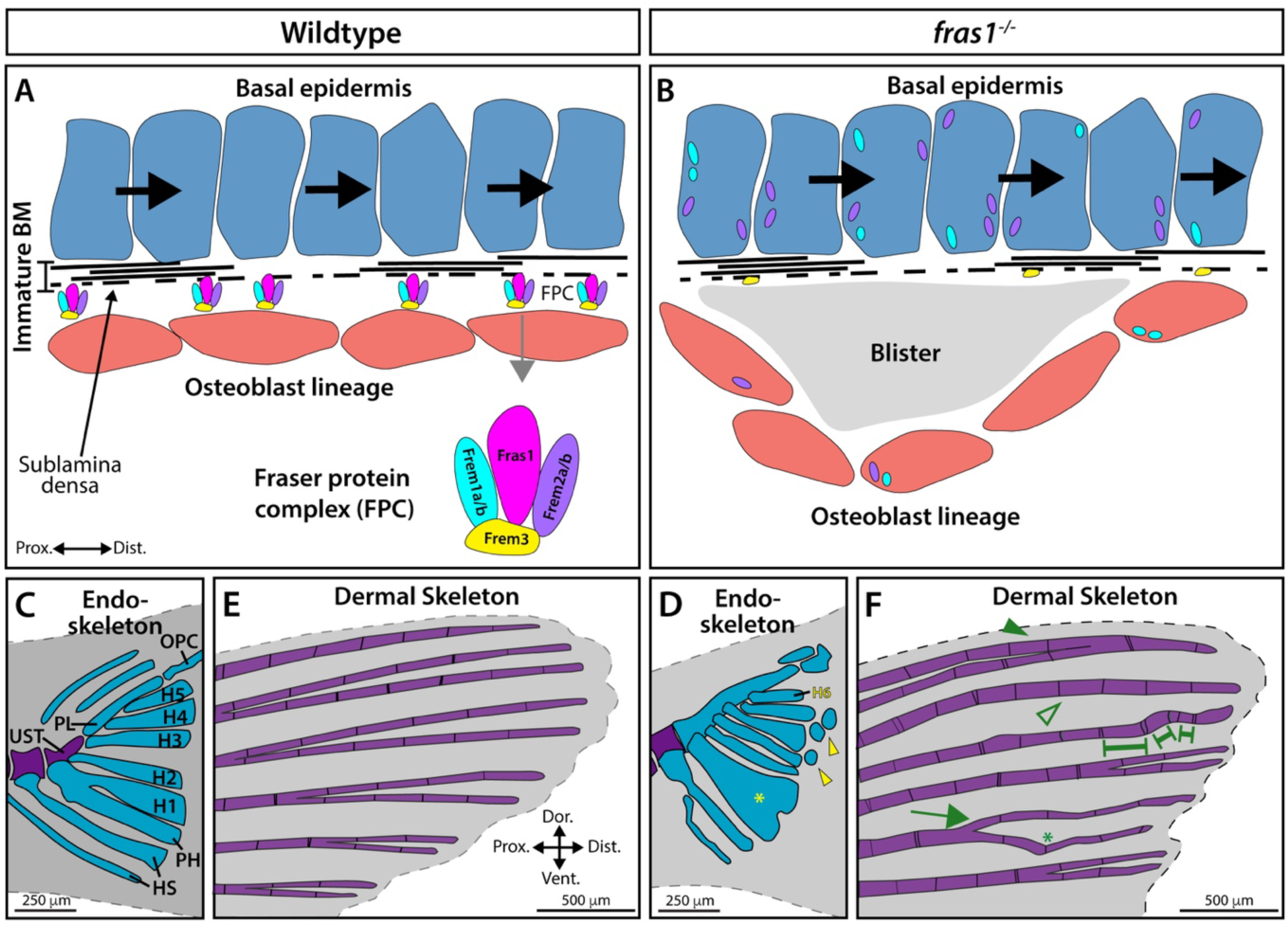
The Fraser Complex promotes growth zone epidermal – mesenchymal tissue connectivity enabling robust fin skeletal development and regeneration. **(A, B)** Schematic of epithelial basal epidermis (blue cells) and mesenchymal pre-osteoblast (salmon cells) tissue organization separated by an immature basement membrane (BM, black dashed lines) in the distal developing or regenerating fin ray. **(A)** The Fraser Complex (FPC) adheres epidermal and mesenchymal tissue layers to support osteoblast organization and therefore skeletal patterning. **(B)** In *fras1^−/−^*mutants, Fraser Complex components cannot translocate to the sublamina densa basement membrane layer and blisters form between tissue layers. **(C)** Wildtype developing caudal endoskeleton with blue and purple representing cartilage and calcified bone, respectively. Abbreviations: UST, urostyle; PL, pleurostyle; OPC, opisthural cartilage; H1-H5, hypurals 1-5; PH, parhypural; HS, hemal spine. **(D)** Example of a *fras1^−/−^* caudal endoskeleton, including ectopic cartilaginous structures (yellow arrowheads), an additional hypural (H6, yellow), fusions between elements (yellow asterisk), and general poor articulation. **(E)** Wildtype caudal fin dermal skeleton (fin rays) anatomy. **(F)** Example of *fras1^−/−^* fin ray patterning defects. Abnormalities include ray fusions (green arrowhead), absent branching (open green arrowhead), uneven joints (green brackets), precocious branching (green arrow), and wavy rays (green asterisk).

Fraser Syndrome occurs in approximately 1:10,000 stillbirths and 1:200,000 live births, suggesting ∼ 5% survival of Fraser Syndrome-afflicted fetuses (Divya et al., 2014; Martínez-Frías et al., 1994; Thomas et al., 1986). Fraser Syndrome patients that survive adolescence can achieve normal life spans (Impallomeni et al., 2006). While *fras1^−/−^* mice are mostly embryonic lethal, they likewise can survive to adulthood depending on genetic background (McGregor et al., 2003; Miller et al., 2013; Vrontou et al., 2003). Similarly, we find 10.4% viability for *fras1^te262^* homozygous zebrafish under careful rearing conditions. These homozygous survivors live over a year under standard care. This phenotypic variation suggests the influence of background genetic modifiers, supported by clutch-specific differences in *fras1^te262^* jaw features (Talbot et al., 2012; Talbot et al., 2016) and the exacerbation of *fras1^te262^* phenotypes by site-independent transgene sequences (Kimmel et al., 2021). Alternatively, characteristic variability across species including humans could arise from the Fraser complex largely conferring robustness to organ development. Fraser complex loss-of-function phenotypes then variably manifest depending on stochastic or environmental perturbations with development often even proceeding normally. Long-term viability through adulthood amongst *fras1* loss-of-function escapers suggests Fraser complex functions are particularly important during organ patterning and morphogenesis and are less essential once organs are established.

Zebrafish enables comparing gene function establishing skeletal pattern during fin development and pattern re-establishment upon fin regeneration. We find Fras1 is not necessary for fin regeneration in general, including outgrowth extent. However, skeletal patterning, including ray branching, defects worsen in regenerated fins lacking Fras1. Therefore, the Fraser Complex both helps establish the overall number and distribution of rays during early fin development and actively supports ray morphogenesis during developmental and regenerative fin outgrowth.

The ray branching morphogenesis defects and mis-aligned pre-osteoblasts in *fras1* mutants tie the Fraser complex to the maintenance of a well-organized environment supporting pre-osteoblast positioning. For Shh/Smo-mediated ray branching, intercellular signaling remains intact but downstream cell behaviors, which could include basal epidermal and pre-osteoblast heterotypic associations and co-movements (Braunstein et al., 2021), likely are perturbed. Frequent blistering distal to the rays in *fras1* mutants suggest the Fraser Complex serves a permissive role sustaining a robustly layered epidermis closely associated with underlying blastema mesenchyme. Similar blistering defects in Fraser complex mutants are observed during development, including for craniofacial patterning (Talbot et al., 2012, 2016). Further, parallel genetic studies in mice describe extensive *in utero* blistering of eye, limb bud, and kidneys that typically become hemorrhagic and are associated with poor outcomes (McGregor et al., 2003; Reviewed in Smyth and Scambler, 2005; Vrontou et al., 2003). Zebrafish fin ray regeneration, including osteoblast-positioning for ray branching, provides an accessible model to understand how Fraser Complex-promoted tissue layer associations enable coordinated epithelial-mesenchymal interactions during tissue morphogenesis.

We find *fras1* is highly enriched in distal basal epidermis of regenerating fins while the *frem* family Fraser Complex subunits are expressed in basal epidermis as well as fibroblast and osteoblast lineage cells of the blastemal mesenchyme. Immunohistochemistry and *in situ* hybridization revealed Fras1 protein and *frem2a/b* transcripts are most highly localized to the distalmost basement membrane separating the intra-ray blastemal mesenchyme from the encasing stratified epidermis. We show that *fras1^−/−^* mutants exhibit diminished Frem2 at the basement membrane but maintain distal basal epidermis intra-cellular expression. Similarly, Frem2 is unable to localize to the basement membrane lamina densa and is instead retained intracellularly in epithelial cells upon *Fras1* loss-of-function in mice (Petrou et al., 2007). Therefore, zebrafish regenerating fins provide a compelling setting to examine mechanisms of Fraser Complex transcription upregulation as well as protein synthesis, processing, and assembly.

The Fraser complex-enriched distal basal epidermis of regenerating fins has an immature basement membrane compared with the thicker and well-defined laminin-containing layer that separates the basal epidermis from mature bone-lining osteoblasts more proximally (Armstrong et al., 2017; Chen et al., 2015). We propose the Fraser complex may serve as a scaffold stabilizing tissue organization during basement membrane formation to enable morphogenetic epithelial-mesenchymal interactions. Extending this idea, the Fraser complex may even template efficient basement membrane assembly. Such general, permissive roles could explain the strikingly varied phenotypes across organs that characterize Fraser Syndrome and animal models thereof. Similarly, a transient function while organs are forming could explain why the Fraser Complex seems more critical during embryonic developmental windows or epimorphic organ regeneration rather than for organ homeostasis (Talbot et al., 2016). Regardless, zebrafish fins provide a new context to explore Fraser Complex mechanisms during epithelial-mesenchymal interactions and, potentially, conserved origins of Fraser Syndrome syndactylism.

## MATERIALS & METHODS

### Zebrafish husbandry and lines

*Danio rerio* zebrafish were housed at the University of Oregon with husbandry as described (Westerfield, 2007) overseen by the Aquatic Animal Care Services (AqACS). Protocols were approved by the Institutional Care and Use Committee (IACUC). The following mutant and transgenic lines were used: *fras1^te262^* line (Carney et al., 2010)*, Tg(−2.4shha:gfpABC)* (Ertzer et al., 2007) and *Tg(ptch2:Kaede)* (Huang et al., 2012).

*fras1^te262^* homozygous mutants were screened by embryonic fin fold blistering at 24 hpf. Larvae were reared in beakers with lower water volume, a reduced concentration of rotifer feed, and health-checked more frequently compared to standard care. At 30 dpf, healthier juveniles were moved to standard housing with no further exceptional care. *fras1^te262^* heterozygote adults were identified by fin clip PCR and subsequent SspI-HF restriction enzyme digest with the following primers:

F: 5’GGAAGATTTTCTTTATTTAGCAGTCTCT
R: 5’ TTGGAACTAGGTCCTCTTTGGTGTGCTATAAAATA

### Skeletal Staining

Live adult fish were stained with Alizarin Red S (LabChem, Zelienople, PA) at 0.01% concentration in fish facility system water for 15 minutes (Bensimon-Brito et al., 2016), followed by 3 × 5 minute washes in fresh system water. Live adult fish were anesthetized in 0.6 mM tricaine (MS222, Syndel) in system water and imaged on a Nikon T*i*-E inverted microscope with a Yokogawa CSU spinning disc confocal unit. Images of all fins (*n* = 5 controls, *n* = 8 *fras1^te262^* mutants) were acquired by tricaine overdose euthanasia of Alizarin-stained fish and subsequent dissection of fins, followed by imaging with dual Brightfield and Widefield fluorescence on a Nikon T*i*-E inverted microscope. Larval fish less than 30 dpf were stained with Alcian Blue (Anatech, Batte Creek, MI) and Alizarin Red S using a modified double-stain protocol (Walker and Kimmel, 2007) whereby the clearing step of 20% glycerol and 0.25% KOH was increased to 4 days with daily solution changes. Fixed skeletal preps were imaged on a Leica M165 FC stereomicroscope.

### Regeneration assay and fin morphometrics

Adult zebrafish were anaesthetized in 0.6 mM tricaine (MS222, Syndel) in system water and fins amputated with a straight-edge razor blade in between the 1^st^ and 2^nd^ procurrent ray, or approximate equivalent when those landmarks were absent in *fras1^−/−^* mutants. Fish immediately were imaged on a Nikon T*i*-E inverted widefield microscope and returned to fresh system water and care. Fish were re-anesthetized at 3, 7, 14 (data not shown), and 28 dpa. *fras1^−/−^* mutants were co-housed with controls at low tank density to facilitate individual tracking (*n* = 4-5 mutants with *n* = 3 controls per tank). Fin skeletal morphometrics were measured using Fiji ImageJ (NIH). Ray lengths were measured from the first discernable proximal joint to the most distal bony segment.

### Immunostaining

Regenerating fins were amputated with a straight razor blade and tissue collected at 3 or 4 dpa, fixed, decalcified, and paraffin embedded as described (Stewart et al., 2014). 7 μm sections were cut using a Leica RM255 microtome. Sections were dehydrated and deparaffinized, rehydrated, and antigen retrieved in 0.05% citranonic anhydride (ImmunoSaver, Electron Microscopy Science) for 5 min on high using a pre-heated pressure cooker. After 5 min, the pressure cooker was turned off and allowed to slowly depressurize for 30 min before the slides were transferred to 1x PBS. Sections were blocked in 1%BSA/1%DMSO/1%NGS/1%FBS in PBS+0.1% Tween-20 (PBST). An anti-FRAS1 human antibody with 76% immunogen identity with zebrafish Fras1 including part of a highly conserved Calx-beta domain (Bethyl Labs, Cat #A305-622A) and anti-FREM2 human antibody with 77% immunogen identity with zebrafish Frem2a/b (Invitrogen PA5-55986) were used. Both primary antibodies were used at 1:250 in blocking solution and incubated overnight at 4°C in a humidified chamber. The next day, sections were washed 3 x 10 min in a high salt solution of 500 mM NaCl in PBST, followed by 1x PBS rinse. Secondary-conjugated AlexaFluor antibodies (Invitrogen) were used at 1:1000 in PBST for 1 hour protected from light. Slides were washed 3 x 5 min in PBS, followed by 1:1500 Hoechst stain (stock 10 mg/mL, Invitrogen) for 5 min, slides washed, and mounted in SlowFade Gold (Invitrogen). Slides were imaged on a Nikon T*i*-E inverted microscope with a Yokogawa CSU-W1 spinning disc confocal attachment.

### In situ hybridization

mRNA transcripts were visualized using the above described 7 μm paraffin sections of regenerating zebrafish fins. mRNA probes for zebrafish *frem2a* and *frem2b* were obtained and visualized using ViewRNA Tissue Assay technology (Invitrogen, #19942) according to manufacturer’s instructions. After probe hybridization and amplification, nuclei were stained with 1:1500 Hoechst stain (Invitrogen), mounted in SlowFade Gold (Invitrogen), and imaged as described above.

### Transcriptomic Analysis

Single Cell RNA-seq expression profiling used a previously assembled and validated dataset comprised of regenerating fin tissue collected at 3- and 7-days post amputation (Lewis et al., 2023). The “choose_cells” function was used to reduce the extended dataset to only include epidermal, fibroblast and osteoblast lineages. Uniform Manifold Approximation and Projection (UMAP) plots detailing the cell-type clustering and expression profiles of Frasier complex genes were generated in Monocle3 using the “Plot_cells” function with otherwise default parameters.

### Shh/Ptch2 reporter line assessment

For Shh ligand expression, caudal fins of adult *shha:GFP* heterozygotes were amputated as described above and imaged with a Nikon T*i*-E inverted widefield fluorescence microscope at 4 (not shown), 8, 14 (not shown), and 30 dpa. Developing *shha:GFP* 6 wpf juveniles were imaged with a Nikon T*i*-E inverted widefield microscope using a Yokogawa CSU-W1 spinning disc attachment. *ptch2:Kaede* adults and juveniles had dorsal portions of their caudal fins photoconverted to red emission with confocal images acquired before photoconversion (all green, not shown), immediately after photoconversion (all red), and 24 hours after conversion (newly produced Kaede is green). Photoconversion used a 20x confocal objective set to 100% 405 λ laser for 1 min (juveniles) or 2 min (regenerating adults). Fish were anesthetized as described for imaging and then returned to standard care.

### Hematoxylin and Eosin Histology

7 μm paraffin sections were collected as described above and deparaffinized with xylenes. Slides were stained with Hematoxylin (Ricca Chemical CAT#3530-16) and Eosin (Sigma-Aldrich REF#HT110132) according to the manufacturer’s protocol with the durations: filtered hematoxylin 1 minute, eosin 4 minutes. Slides were then dehydrated and mounted using Permount Mounting Media (Electron Microscopy Sciences CAT#17986). Brightfield images were acquired on a Leica DM4000B widefield microscope and confocal Eosin images were taken using a Nikon T*i*-E with a Yokogawa W1-CSU confocal unit.

### Statistical Analyses

Fin morphometric data was collected using Fiji ImageJ (NIH) and organized in Microsoft Excel. Fin width was measured from the most proximal distinguishable joint segment. Ray length was measured starting at the same location. Number of principal rays was quantified prior to ray branching, if present. Statistical analyses were performed and graphs generated using GraphPad Prism v9.

**Table 1.** Fin morphometrics associated with Figure 1. Quantification of principal ray number, procurrent ray number, and fraction of branched principal rays for the adult caudal fins of indicated genotype and shown in denoted Fig. 1 panels.

| **Fig. 1 Panel** | **# Principal Rays** | **# Procurrent Rays** | **Branched/Total Principal Rays** |
| --- | --- | --- | --- |
| **B (*fras1<sup>+/-</sup>*)** | **18** | **6** | **16/18 (88.9%)** |
| **C (*fras1<sup>-/-</sup>*)** | **14** | **5** | **11/14 (78.6%)** |
| **D (*fras1<sup>-/-</sup>*)** | **11** | **4** | **6/11 (54.5%)** |
| **E (*fras1<sup>-/-</sup>*)** | **12** | **3** | **5/12 (41.7%)** |
| **F (*fras1<sup>-/-</sup>*)** | **4** | **N/A** | **1/4 (25.0%)** |

## ACKNOWLEDGEMENTS

We thank the University of Oregon AqACS Facility for zebrafish care; the University of Oregon zebrafish community for support; and C. Kimmel, T. Desvignes, and the Stankunas lab for input.

## FOOTNOTES

### Author contributions

A.E.R. and K.S. designed experiments with input from S.S.; A.E.R., S.H., and V.M.L. performed experiments; A.E.R. and K.S. prepared and wrote the manuscript.

### Data and material availability

Requests for materials should be addressed to K. S.

---

**Figure 1 Supplement 1.**
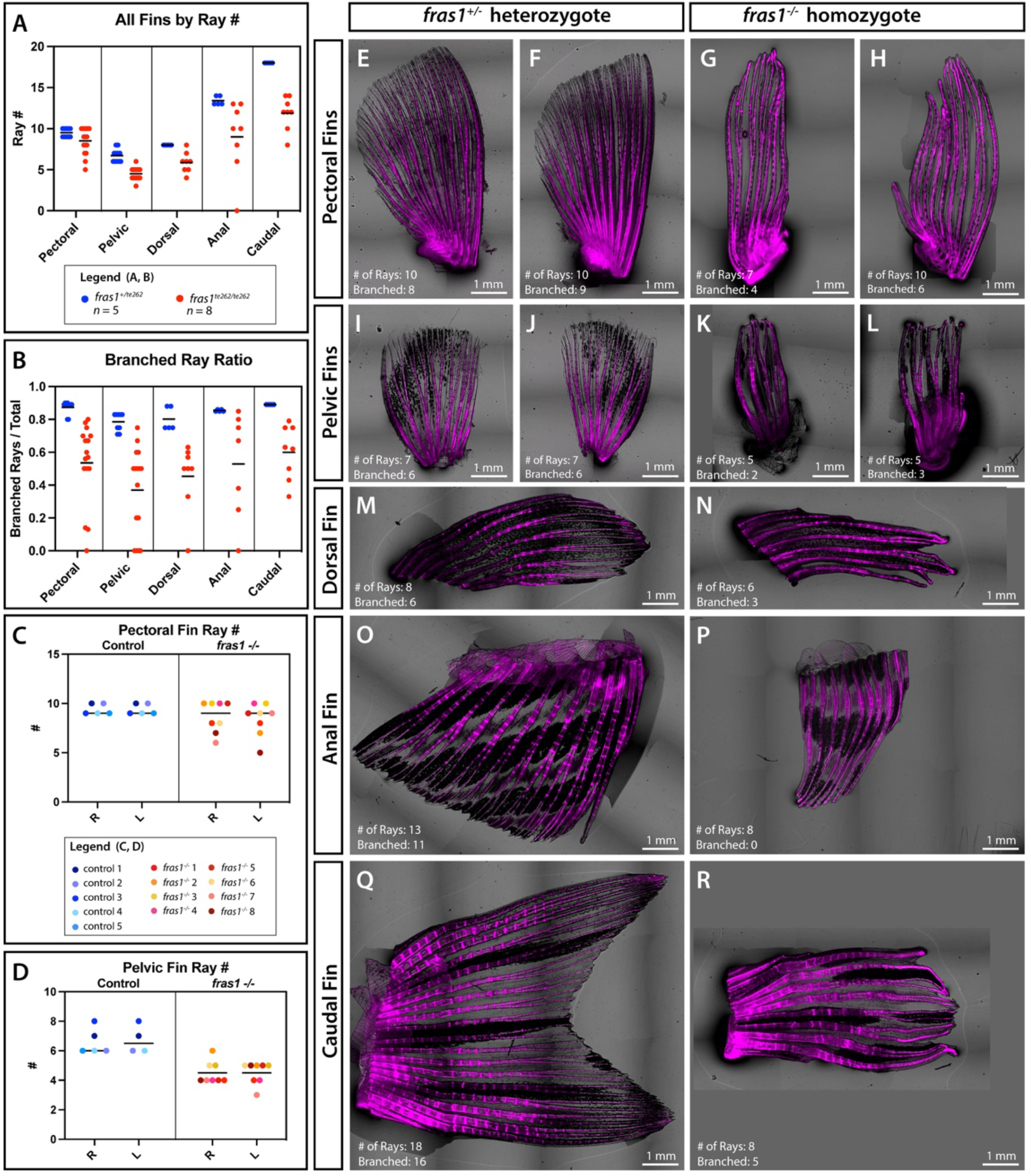
Variable expressivity of *fras1^−/−^* skeletal patterning abnormalities across all fins. **(A)** Fin ray scoring for all 7 zebrafish adult fins for *n* = 5 *fras1^+/−^* heterozygote controls and *n* = 8 *fras1^−/−^* homozygote mutants. The paired pectoral and pelvic fins have two data points per fish. **(B)** Ratio of branched fin rays over total fin rays per fin. The paired pectoral and pelvic fins have two data points per fish. **(C)** Pectoral fin ray number and **(D)** Pelvic fin ray number for right (R) and left (L) fins with data points for individual control and *fras1^−/−^* fish in matching blue or red spectrum colors, respectively. Means are shown. **(E-R)** Representative Alizarin Red-stained adult caudal fins overlayed with brightfield for **(E, F, I, J, M, O, Q)** controls and **(G, H, K, L, N, P, R)** *fras1*^−/−^ mutants. **(E-H)** Pectoral and **(I-L)** pelvic fins are from the same animal to highlight left-right asymmetry in *fras1^−/−^* mutants. Number of rays, branched rays, and scale bars are as marked.

**Figure 4 Supplement 1.**
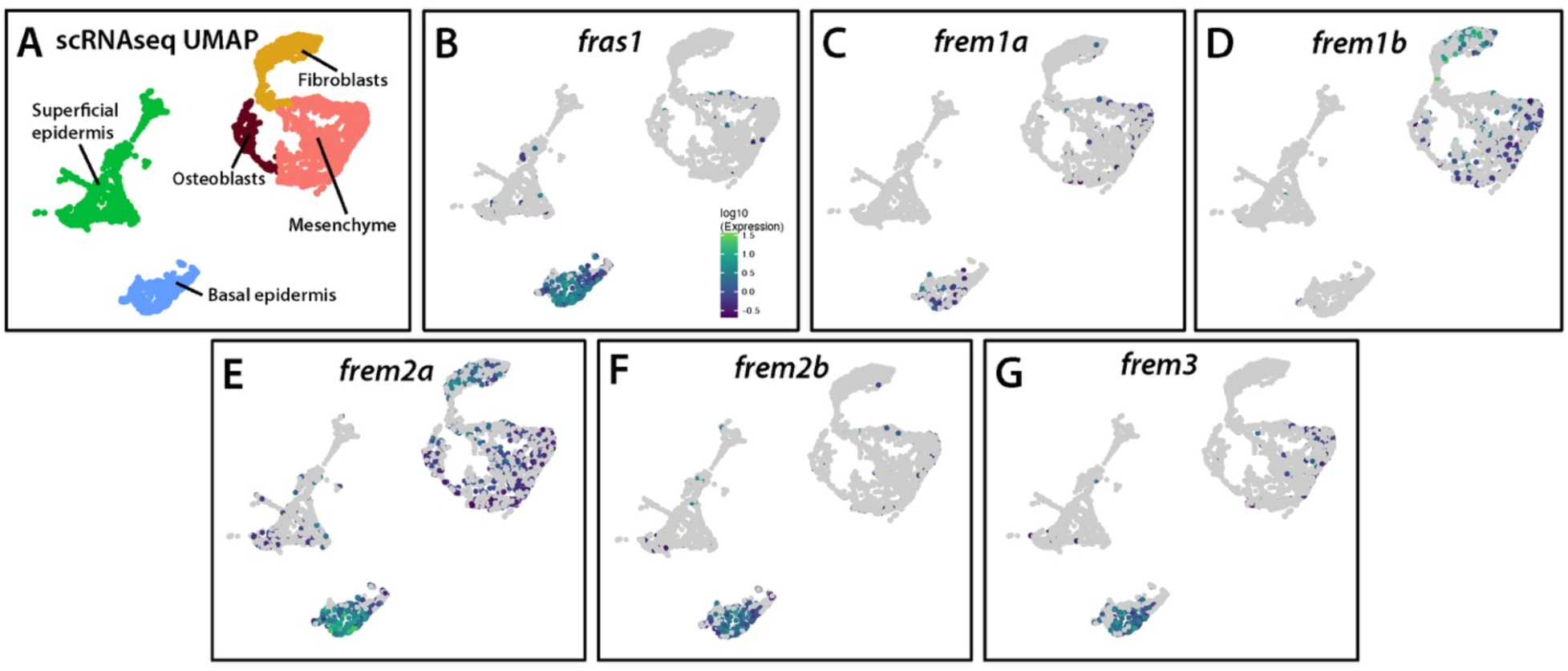
Single cell-resolved gene expression profiles show Fraser Complex transcripts are basal epidermal-enriched during fin regeneration. **(A)** UMAP cluster analysis from single cell transcriptomes of 3- and 7-day post amputation (dpa) zebrafish fin tissue identifying unique clusters for basal epidermis (blue), superficial epidermis (green), and mixed blastemal mesenchyme / osteoblasts / fibroblasts (dark red, pink, yellow). **(B-G)** Individual UMAP plots for Fraser Complex components *fras1*, *frem1a*, *frem1b*, *frem2a*, *frem2b*, and *frem3*. *fras1* is the most highly enriched basal epidermal marker of all captured transcripts.

**Figure 6 Supplement 1.**
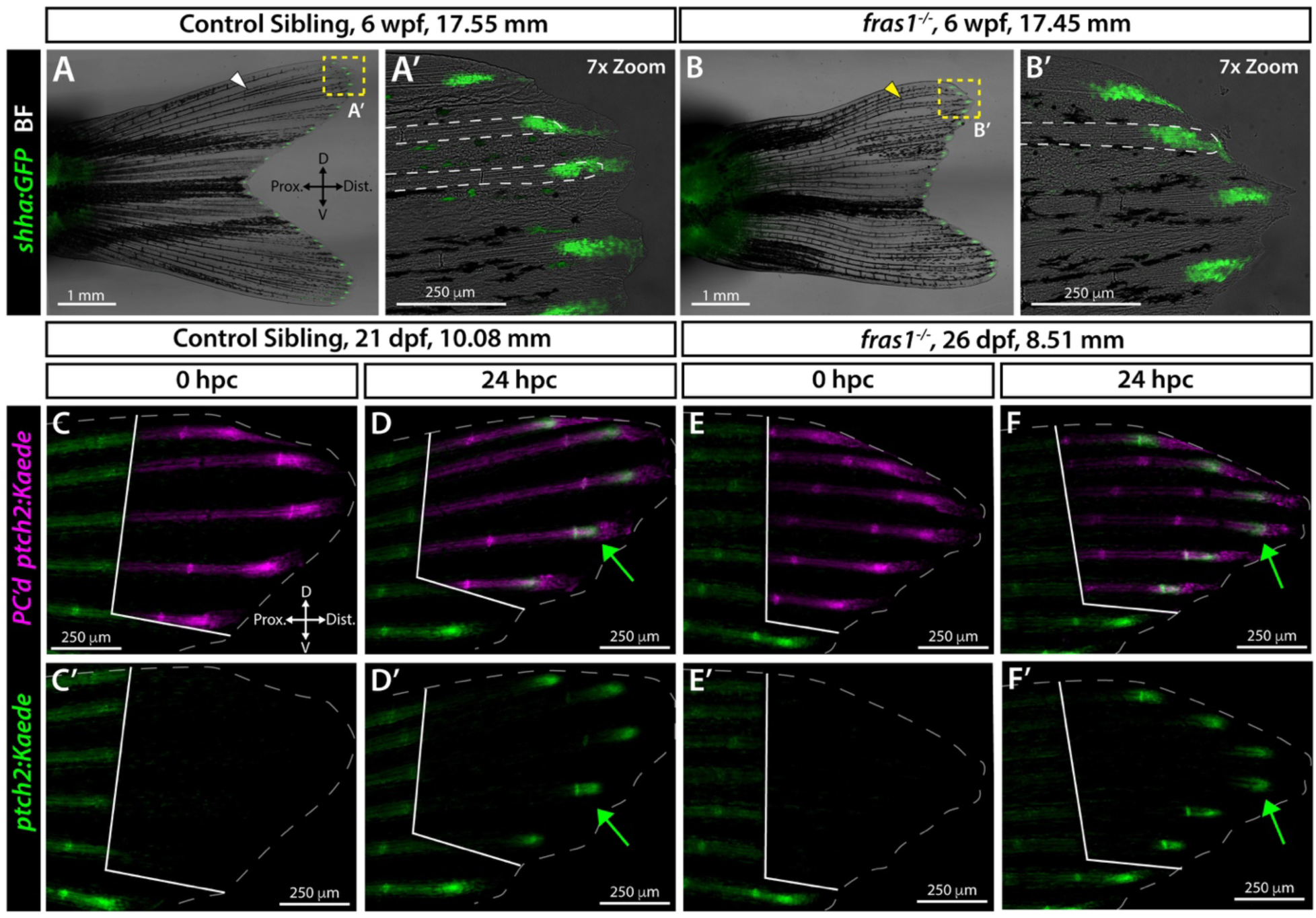
Loss of Fras1 does not impair Shh/Smo signaling during fin development. **(A, B)** Whole mount caudal fin images of representative **(A)** *shha:GFP* control (*n* = 4) and **(B)** *shha:GFP; fras1^−/−^* (*n* = 6) siblings at six weeks post fertilization (wpf). **(A’, B’)** Zoom insets of the dorsal tip of the caudal fins. Dorsal ray 2 (dashed white outline) is branched in **(A’)** but unbranched in **(B’)**. **(C-F’)** Confocal fluorescence images of the dorsal lobe of developing caudal fins from a *ptch2:Kaede* control and stage-matched *fras1^−/−^*mutant sibling (*n* = 3 controls, *n* = 8 mutants). **(C, E)** White outlined regions show complete conversion of Kaede from green to red emission at 0 hours-post-conversion (hpc) following UV light exposure (single channel panels in **C’, E’**). **(D, F)** At 24 hpc, the same fish express small domains of green Kaede protein (green arrows) newly produced in the photoconverted region (white lines), indicating active Shh/Smo signaling. Dashed gray lines indicate fin boundaries.

